# Chimeric spike mRNA vaccines protect against Sarbecoviru*s* challenge in mice

**DOI:** 10.1101/2021.03.11.434872

**Authors:** David R. Martinez, Alexandra Schäfer, Sarah R. Leist, Gabriela De la Cruz, Ande West, Elena N. Atochina-Vasserman, Lisa C. Lindesmith, Norbert Pardi, Robert Parks, Maggie Barr, Dapeng Li, Boyd Yount, Kevin O. Saunders, Drew Weissman, Barton F. Haynes, Stephanie A. Montgomery, Ralph S. Baric

**Author notes:** Corresponding authors. David R. Martinez, D.R.M., Ralph S. Baric, R.S.B.

## Abstract

The emergence of SARS-CoV in 2003 and SARS-CoV-2 in 2019 highlights the need to develop universal vaccination strategies against the broader *Sarbecovirus* subgenus. Using chimeric spike designs, we demonstrate protection against challenge from SARS-CoV, SARS-CoV-2, SARS-CoV-2 B.1.351, bat CoV (Bt-CoV) RsSHC014, and a heterologous Bt-CoV WIV-1 in vulnerable aged mice. Chimeric spike mRNAs induced high levels of broadly protective neutralizing antibodies against high-risk Sarbecoviruses. In contrast, SARS-CoV-2 mRNA vaccination not only showed a marked reduction in neutralizing titers against heterologous Sarbecoviruses, but SARS-CoV and WIV-1 challenge in mice resulted in breakthrough infection. Chimeric spike mRNA vaccines efficiently neutralized D614G, UK B.1.1.7., mink cluster five, and the South African B.1.351 variant of concern. Thus, multiplexed-chimeric spikes can prevent SARS-like zoonotic coronavirus infections with pandemic potential.

**Sentence:** Chimerized RBD, NTD, and S2 spike mRNA-LNPs protect mice against epidemic, zoonotic, and pandemic SARS-like viruses

## Introduction

A novel severe acute respiratory syndrome coronavirus (SARS-CoV) emerged in 2003 and caused more than 8,000 infections and ~800 deaths worldwide (*1*). Less than a decade later, the Middle East Respiratory Syndrome (MERS-CoV) coronavirus emerged in Saudi Arabia in 2012 (*2*), with multiple outbreaks that have resulted in at least ~2,600 cases and 900 deaths (*3*). In December 2019, another novel human SARS-like virus from the genus *Betacoronavirus* and subgenus *Sarbecovirus* emerged in Wuhan China, designated SARS-CoV-2, causing the COVID-19 pandemic (*4, 5*).

Bats are known reservoirs of SARS-like coronaviruses (CoVs) and harbor high-risk “pre-emergent” SARS-like variant strains, such as WIV-1-CoV and RsSHC014-CoV, which are able to use human ACE2 receptors for entry, replicate efficiently in primary airway epithelial cells, and in mice, and may escape existing countermeasures (*6–12*) Given the high pandemic potential of zoonotic and epidemic Sarbecoviruses (*12*), the development of countermeasures, such as broadly effective vaccines, antibodies and drugs is a global health priority (*13–16*).

Sarbecovirus spike proteins have immunogenic domains: the receptor binding domain (RBD), the N-terminal domain (NTD), and the subunit 2 (S2) (*17–20*). RBD, NTD, and S2 are a target for neutralizing antibodies elicited in the context of natural SARS-CoV-2 and MERS-CoV infections (*17, 20–24*). In fact, passive immunization with SARS-CoV-2 NTD-specific antibodies protect naïve mice from challenge, demonstrating that the NTD is a target of protective immunity (*18, 24, 25*). However, it remains unclear if vaccine-elicited neutralizing antibodies can protect against *in vivo* challenge with heterologous epidemic and bat coronaviruses. Here, we generated nucleoside-modified mRNA-lipid nanoparticle (LNP) vaccines expressing chimeric spikes containing admixtures of RBD and NTD domains from zoonotic, epidemic, and pandemic CoVs and examined their efficacy against homologous and heterologous Sarbecovirus challenge in aged mice.

## Results

### Design and expression of chimeric spike constructs to cover pandemic and zoonotic SARS-related coronaviruses

Sarbecoviruses exhibit considerable genetic diversity (Fig. 1A) and SARS-like bat CoVs (Bt-CoVs) are recognized threats to human health (*8, 12*). Harnessing the modular structure of CoV spikes (*26*), we designed chimeric spikes by admixture of divergent clade I-III Sarbecovirus NTD, RBD, and S2 domains into “bivalent” and “trivalent” vaccine immunogens that have the potential to elicit broad protective antibody responses against distant strains (e.g., *Sarbecovirus*). The approach is designed to maximize immune breadth in monovalent and multiplexed formulations. We designed four sets of chimeric spike constructs that contained admixtures of the RBD and/or NTD, and S2 neutralizing domains from various Sarbecoviruses. Chimera 1 included the NTD from clade II Bt-CoV Hong Kong University 3-1 (HKU3-1), the clade I SARS-CoV RBD, and the clade III SARS-CoV-2 S2 (Fig. 1B). Chimera 2 included SARS-CoV-2 RBD and SARS-CoV NTD and S2 domains (*16*). Chimera 3 included the SARS-CoV RBD, and SARS-CoV-2 NTD and S2, while chimera 4 included the RsSHC014 RBD, and SARS-CoV-2 NTD and S2. We also generated a monovalent SARS-CoV-2 spike furin knock out (KO) vaccine, partially phenocopying the Moderna and Pfizer mRNA vaccines in human use, and a negative control norovirus GII capsid vaccine (Fig. 1B, 1C). We generated these chimeric spikes and control spikes as lipid nanoparticle-encapsulated, nucleoside-modified mRNA vaccines with LNP adjuvants (mRNA-LNP) as described previously (*27*). This mRNA LNP stimulate robust T follicular helper cell activity, germinal center B cell responses, and durable long-lived plasma cells and memory B cell responses (*20, 28*). We verified their chimeric spike expression in HEK cells (Fig. S1B). To confirm that scrambled coronavirus spikes are biologically functional, we also designed and recovered several high titer recombinant live viruses of RsSHC014/SARS-CoV-2 S1, NTD, RBD and S2 domain chimeras that included deletions in non-essential, accessory ORF7&8 and that encoded nanoluciferase (Fig. S1C).

**Figure 1.**
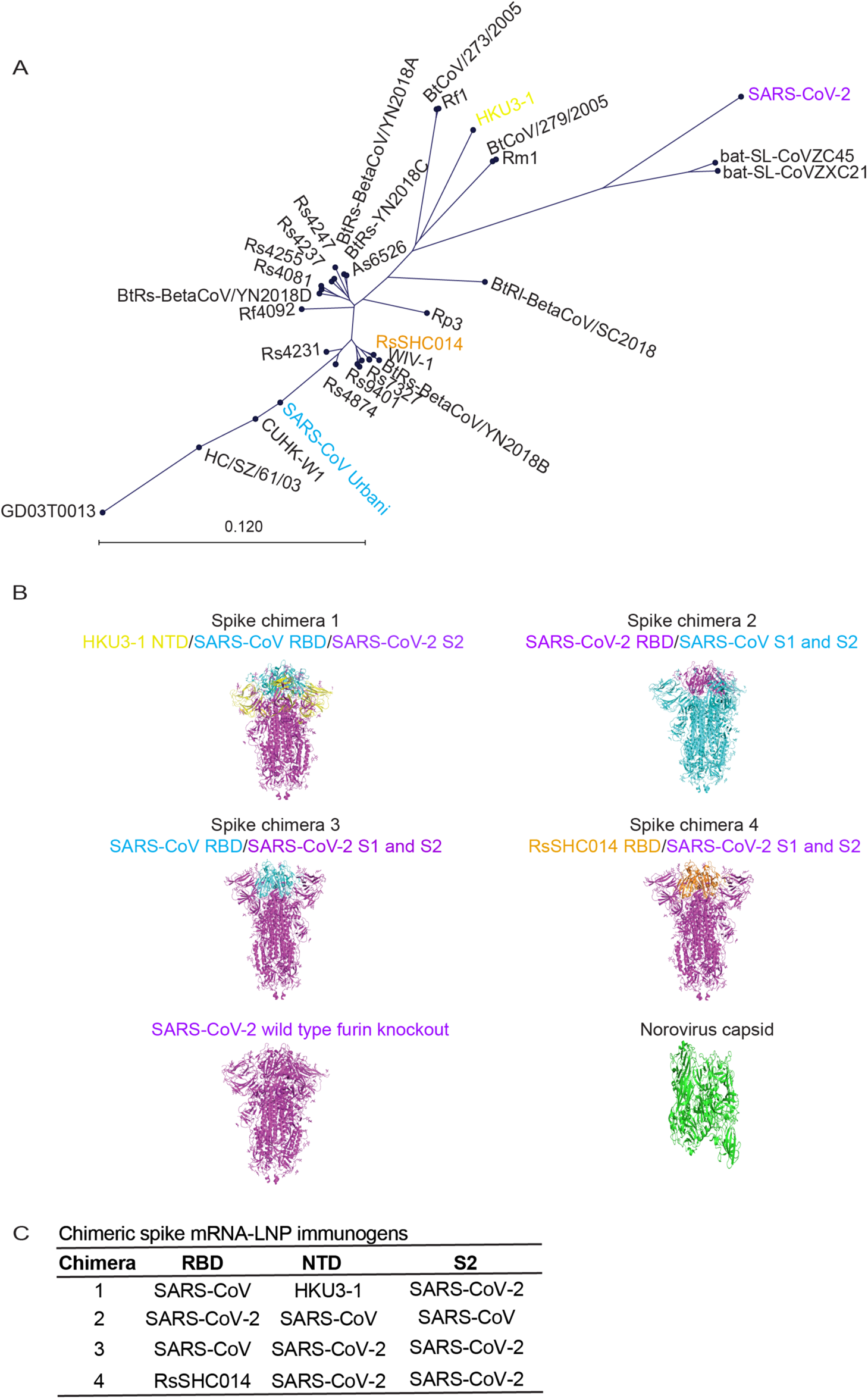
Genetic design of chimeric Sarbecovirus spike vaccines. (**A**) Genetic diversity of pandemic and bat zoonotic coronaviruses. SARS-CoV is shown in light blue, RsSHC014 is shown in purple, and SARS-CoV-2 is shown in red. (**B**) Spike chimera 1 includes the NTD from HKU3-1, the RBD from SARS-CoV, and the rest of the spike from SARS-CoV-2. Spike chimera 2 includes the RBD from SARS-CoV-2 and the NTD and S2 from SARS-CoV. Spike chimera 3 includes the RBD from SARS-CoV and the NTD and S2 SARS-CoV-2. Spike chimera 4 includes the RBD from RsSHC014 and the rest of the spike from SARS-CoV-2. SARS-CoV-2 furin KO spike vaccine and is the norovirus capsid vaccine. (**C**) Table summary of chimeric spike constructs.

### Immunogenicity of mRNAs expressing chimeric spike constructs against coronaviruses

We next sought to determine if simultaneous immunization with mRNA-LNP expressing the chimeric spikes of diverse Sarbecoviruses was a feasible strategy to elicit broad binding and neutralizing antibodies. We immunized aged mice with the chimeric spikes formulated to induce type-specific and/or cross-reactive responses against multiple divergent clade I-III Sarbecoviruses, a SARS-CoV-2 furin KO spike, and a GII.4 norovirus capsid negative control. Group 1 was primed and boosted with chimeric spikes 1, 2, 3, and 4 (Fig. S1A). Group 2 was primed with chimeric spikes 1 and 2 and boosted with chimeric spikes 3 and 4 (Fig. S1A). Group 3 was primed and boosted with chimeric spike 4 (Fig. S1A). Group 4 was primed and boosted with the monovalent SARS-CoV-2 furin knockout spike (Fig. S1A). Finally, group 5 was primed and boosted with a norovirus capsid GII.4 Sydney 2011 strain (Fig. S1A). We then examined the binding antibody responses by ELISA against a diverse panel of CoV spike proteins that included epidemic, pandemic, and zoonotic coronaviruses.

Mice in groups 1 and 2 generated the highest magnitude responses to SARS-CoV Toronto Canada isolate (Tor2), RsSHC014, and HKU3-1 spike compared to group 4 (Fig 2A, 2G, and 2H). While mice in group 2 generated lower magnitude binding responses to both SARS-CoV-2 RBD (Fig. 2C) and SARS-CoV-2 NTD (Fig 2D), mice in group 1 generated similar magnitude binding antibodies to SARS-CoV-2 D614G compared to mice immunized with the SARS-CoV-2 furin KO spike mRNA-LNP (Fig 2B). Mice in groups 1 and 2 generated similar magnitude binding antibody responses against SARS-CoV-2 D614G, Pangolin GXP4L, and RaTG13 spikes (Fig. 2B, 2E, and 2F) compared to mice from group 4. Mice in group 1 and group 4 elicited high magnitude levels of hACE2 blocking responses, as compared to groups 2 and 3 (Fig. 2J). As binding antibody responses post boost mirrored the trend of the post prime responses, it is likely that the second dose is boosting immunity to the vaccine antigens in the prime (Fig. 2). Finally, we did not observe cross-binding antibodies against common-cold CoV spike antigens from HCoV-HKU1, HCoV-NL63, and HCoV-229E in most of the vaccine groups (Fig. S2A-2D), but we did observe low binding levels against more distant group 2C MERS-CoV (Fig. 2I) and other Betacoronaviruses like group 2A HCoV-OC43 in vaccine groups 1 and 2 (Fig. S2B). These results suggest that chimeric spike mRNA vaccines elicit broader and higher magnitude binding responses against pandemic and bat SARS-like viruses compared to monovalent SARS-CoV-2 spike mRNA-LNP vaccines.

**Figure 2.**
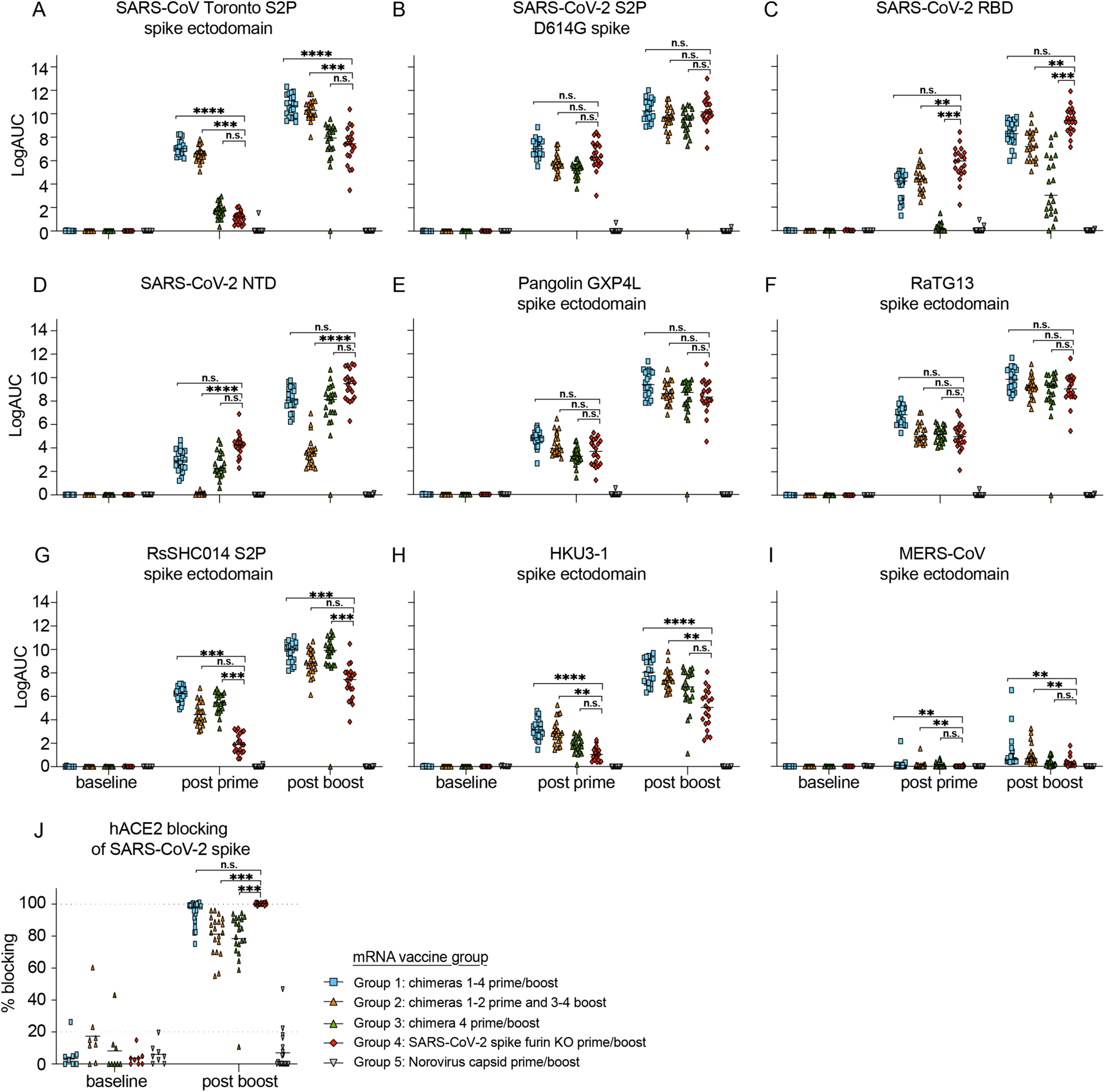
Human pathogenic coronavirus spike binding and hACE2-blocking responses in chimeric and monovalent SARS-CoV-2 spike-vaccinated mice. Serum antibody ELISA binding responses were measured in the five different vaccination groups. Pre-immunization, post prime, and post-boost binding responses were evaluated against Sarbecoviruses, MERS-CoV, and common-cold CoV antigens including: (**A**) SARS-CoV Toronto Canada (Tor2) S2P, (**B**) SARS-CoV-2 S2P D614G, (**C**) SARS-CoV-2 RBD, (**D**) SARS-CoV-2 NTD, (**E**) Pangolin GXP4L spike, (**F**) RaTG13 spike, (**G**) RsSHC014 S2P spike, (**H**) HKU3-1 spike, (**I**) MERS-CoV spike, (**J**) hACE2 blocking responses against SARS-CoV-2 spike in the distinct immunization groups. Blue squares represent mice from group 1, orange triangles represent mice from group 2, green triangles represent mice from group 3, red rhombuses represent mice from group 4, and upside-down triangle represent mice from group 5. Statistical significance for the binding and blocking responses is reported from a Kruskal-Wallis test after Dunnett’s multiple comparison correction. *p < 0.05, **p < 0.01, ***p < 0.001, and ****p < 0.0001.

### Neutralizing antibody responses against live Sarbecoviruses and variants of concern

We then examined the neutralizing antibody responses against SARS-CoV, Bt-CoV RsSHC014, Bt-CoV WIV-1, and SARS-CoV-2 and variants of concern using live viruses as previously described (Fig 3A-3D) (*29*). Group 4 SARS-CoV-2 S mRNA vaccinated animals mounted a robust response against SARS-CoV-2, however responses against SARS-CoV, RsSHC014, and WIV-1 were 18-, >500- or 116-fold more resistant, respectively (Fig 3A-3D and Fig. S3G-H). In contrast, aged mice in group 2 showed a 42- and 2-fold increase in neutralizing titer against SARS-CoV and WIV1, and less than 1-fold decrease against RsSHC014 relative to SARS-CoV-2 neutralizing titers (Fig 3A-3D and Fig. S3C-D). Mice in group 3 elicited 3- and 7-fold higher neutralizing titers against SARS-CoV and RsSHC014 yet showed a 3-fold reduction in WIV-1 neutralizing titers relative to SARS-CoV-2 (Fig 3A-3D and Fig. S3E-F). Finally, mice in group 1 generated the most balanced and highest neutralizing titers that were 13- and 1.2-fold higher against SARS-CoV and WIV-1 and less than 1-fold lower against RsSHC014 relative to the SARS-CoV-2 neutralizing titers (Fig 3A-3D and Fig. S3A-B). The serum of mice from groups 1 and 4 neutralized the dominant D614G variant with similar potency as the wild type D614 non-predominant variant, and both groups had similar neutralizing antibody responses against the U.K. B.1.1.7 and the mink cluster 5 variants as compared to the D614G variant (Fig. 3E 3F). Despite the significant but small reduction in neutralizing activity against the B.1.351 variant of concern (VOC), we did not observe a complete ablation in neutralizing activity in either group. Mice from groups 1 and 2 elicited lower binding and neutralizing responses to SARS-CoV-2 compared to group 4 perhaps reflecting a lower amount of mRNA vaccine incorporated into multiplexed formulations, whereas the monomorphic vaccines may drive a more focused B cell responses to SARS-CoV-2 whereas chimeric spike antigens lead to more breadth against distant Sarbecoviruses. Thus, both monovalent SARS-CoV-2 vaccines and multiplexed chimeric spikes elicit neutralizing antibodies against newly emerged SARS-CoV-2 variants and multiplexed chimeric spike vaccines outperform the monovalent SARS-CoV-2 vaccines in terms of breadth of potency against multiclade Sarbecoviruses.

**Figure 3.**
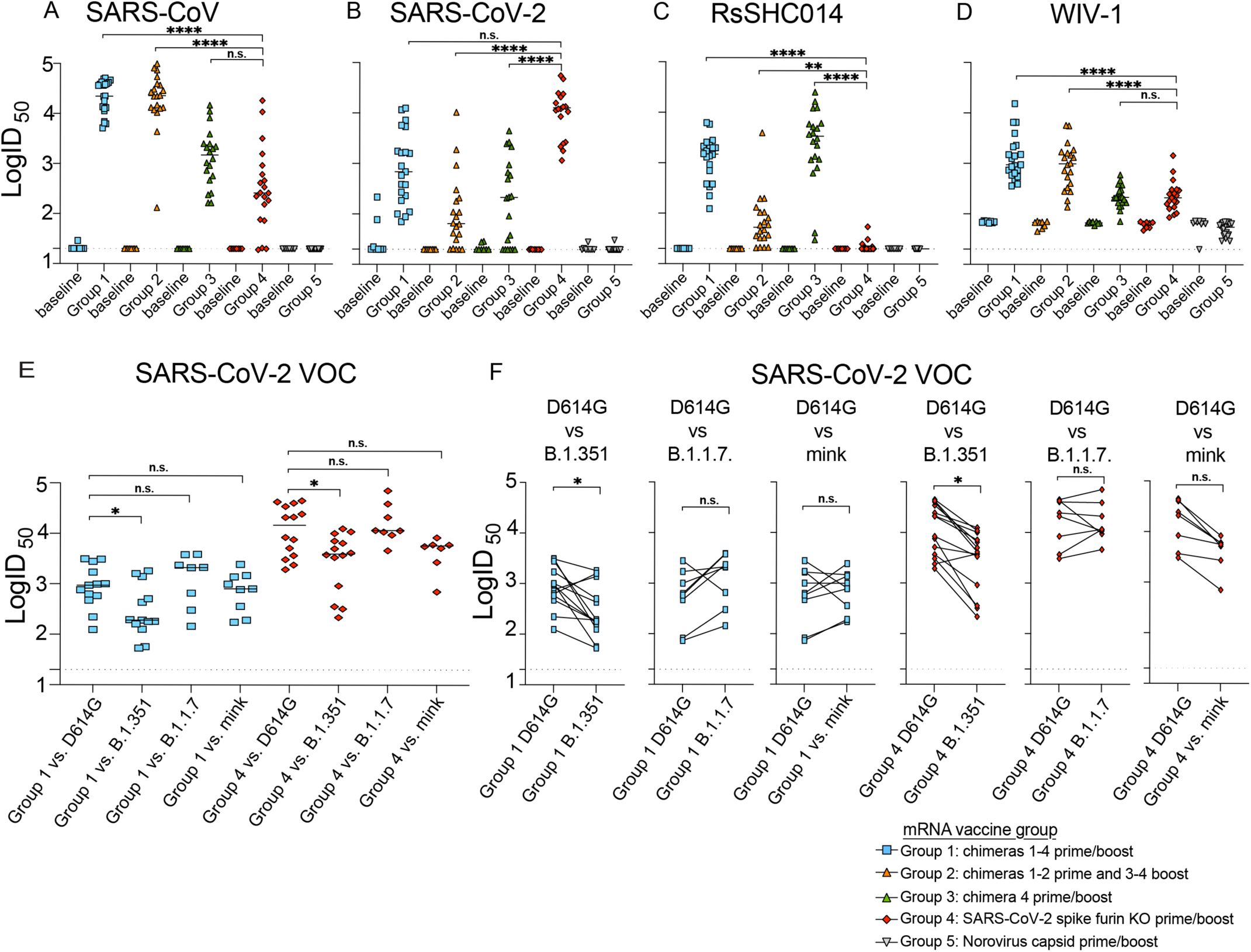
Live Sarbecovirus neutralizing antibody responses in vaccinated mice. Neutralizing antibody responses in mice from the five different vaccination groups were measured using nanoluciferase-expressing recombinant viruses. (**A**) SARS-CoV neutralizing antibody responses from baseline and post boost in the distinct vaccine groups. (**B**) SARS-CoV-2 neutralizing antibody responses from baseline and post boost. (**C**) RsSHC014 neutralizing antibody responses from baseline and post boost. (**D**) WIV-1 neutralizing antibody responses from baseline and post boost. (**E**) The neutralization activity in groups 1 and 4 against SARS-CoV-2 D614G, South African B.1.351, U.K. B.1.1.7, and mink variants (**F**) Neutralization comparison of SARS-CoV-2 D614G vs. South African B.1.351, vs. U.K. B1.1.7, and mink variants. Statistical significance for the live-virus neutralizing antibody responses is reported from a Kruskal-Wallis test after Dunnett’s multiple comparison correction. *p < 0.05, **p < 0.01, ***p < 0.001, and ****p < 0.0001.

### *In vivo* protection against heterologous Sarbecovirus challenge

To assess the ability of the mRNA-LNP vaccines to mediate protection against previously epidemic SARS-CoV, pandemic SARS-CoV-2, and Bt-CoVs, we challenged the different groups and observed the mice for signs of clinical disease. Mice from group 1 or group 2 were completely protected from weight loss, lower, and upper airway virus replication as measured by infectious virus plaque assays following 2003 SARS-CoV mouse-adapted (MA15) challenge (Fig. 4A, 4B and 4C). Similarly, these two vaccine groups were also protected against SARS-CoV-2 mouse-adapted (MA10) challenge. In contrast, group 3 showed some protection against SARS-CoV MA15 induced weight loss, but not against viral replication in the lung or nasal turbinates. Group 3 was fully protected against SARS-CoV-2 MA10 challenge. In contrast, group 5 vaccinated mice developed severe disease including mortality in both SARS-CoV MA15 and SARS-CoV-2 MA10 infections (Fig. S5B, S5C). Monovalent SARS-CoV-2 mRNA vaccines were highly efficacious against SARS-CoV-2 MA10 challenge but failed to protect against SARS-CoV MA15-induced weight loss, and replication in the lower and upper respiratory tract (Fig. 4A, 4B, and 4C), suggesting that SARS-CoV-2 mRNA-LNP vaccines are not likely to protect against future SARS-CoV emergence events. Mice from groups 1-4 were completely protected from weight loss and lower airway SARS-CoV-2 MA10 replication (Fig. 4D, 4E, and 4F). Using both a Bt-CoV RsSHC014 full-length virus and a more virulent RsSHC014-MA15 chimera in mice, we also demonstrated protection in groups 1-3 against RsSHC014 replication in the lung and nasal turbinates (Fig. S4) but not in mice that received the SARS-CoV-2 mRNA vaccine. Group 5 control mice challenged with RsSHC014-MA15 developed disease including mortality (Fig. S5D). Group 3 mice, which received a SARS-CoV-2 NTD/RsSHC014 RBD/S2, was fully protected against both SARS-CoV-2 and RsSHC014 challenge whereas group 4 was not, demonstrating that a single NTD and RBD chimeric spike can protect against more than one virus compared to a monomorphic spike.

**Figure 4.**
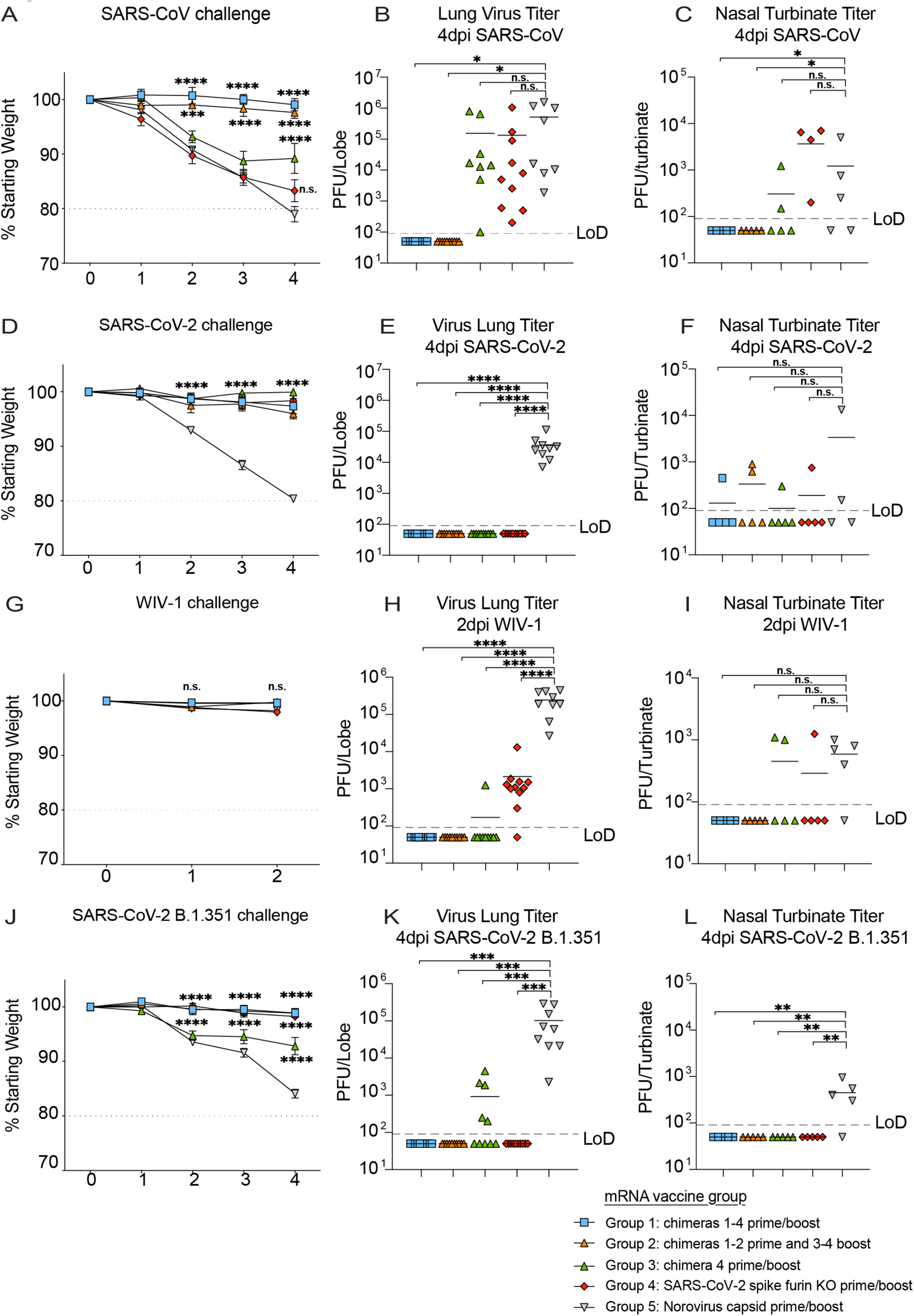
*In vivo* protection against Sarbecovirus challenge after mRNA-LNP vaccination. (**A**) Percent starting weight from the different vaccine groups of mice challenged with SARS-CoV MA15. (**B**) SARS-CoV MA15 lung viral titers in mice from the distinct vaccine groups. (**C**) SARS-CoV MA15 nasal turbinate titers. (**D**) Percent starting weight from the different vaccine groups of mice challenged with SARS-CoV-2 MA10. (**E**) SARS-CoV-2 MA10 lung viral titers in mice from the distinct vaccine groups. (**F**) SARS-CoV-2 MA10 nasal turbinate titers. (**G**) Percent starting weight from the different vaccine groups of mice challenged with WIV-1. (**H**) WIV-1 lung viral titers in mice from the distinct vaccine groups. (**I**) WIV-1 nasal turbinate titers. (**J**) Percent starting weight from the different vaccine groups of mice challenged with SARS-CoV-2 B.1.351. (**K**) SARS-CoV-2 B.1.351 lung viral titers in mice from the distinct vaccine groups. (**L**) SARS-CoV-2 B.1.351 nasal turbinate titers. Figure legend at the bottom right depicts the vaccines utilized in the different groups. Statistical significance for weight loss is reported from a two-way ANOVA after Dunnett’s multiple comparison correction. For lung and nasal turbinate titers, statistical significance is reported from a one-way ANOVA after Tukey’s multiple comparison correction. *p < 0.05, **p < 0.01, ***p < 0.001, and ****p < 0.0001.

We then performed a heterologous challenge experiment with the bat pre-emergent WIV-1-CoV (*9*). Consistent with the protection observed against SARS-CoV, mice from groups 1 and 2 were fully protected against heterologous WIV-1 challenge whereas mice that received the SARS-CoV-2 mRNA vaccine breakthrough replication in the lung (Fig. 5G, 5H, and 5I). We also challenged with a virulent form of SARS-CoV-2 VOC B.1.351, which contains deletions in the NTD and mutations in the RBD, and observed full protection in vaccine groups 1, 2, and 4 compared to controls, whereas breakthrough replication was observed in group 3, further underlining the importance of the NTD in vaccine-mediated protection (Fig. 5J, 5K, and 5L) and its inclusion in universal vaccination strategies. Moreover, the SARS-CoV-2 mRNA vaccine protected against SARS-CoV-2 B.1.351 challenge in aged mice despite a reduction in the neutralizing activity against this VOC.

**Figure 5.**
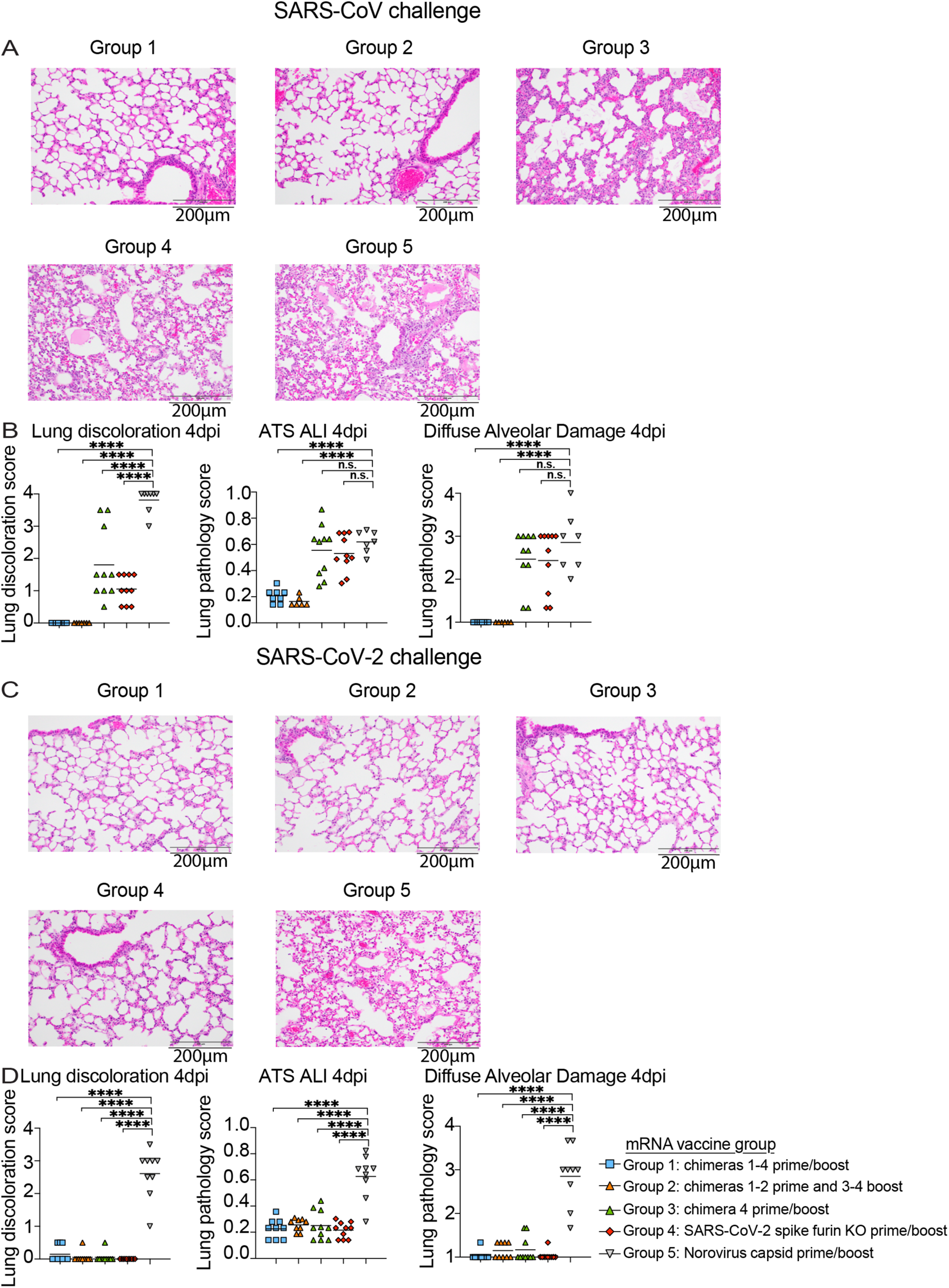
Lung pathology in vaccinated mice after SARS-CoV and SARS-CoV-2 challenge. (**A**) Hematoxylin and eosin 4 days post infection lung analysis of SARS-CoV MA15 challenged mice from the different groups: group 1: chimeras 1-4 prime and boost, group 2: chimeras 1-2 prime and 3-4, group 3: chimera 4 prime and boost, SARS-CoV-2 furin KO prime and boost, and norovirus capsid prime and boost. (**B**) Lung pathology quantitation in SARS-CoV MA15 challenged mice from the different groups. Macroscopic lung discoloration score, microscopic acute lung injury (ALI) score, and diffuse alveolar damage (DAD) in day 4 post infection lung tissues are shown. (**C**) Hematoxylin and eosin 4 days post infection lung analysis of SARS-CoV-2 MA10 challenged mice from the different groups. (**D**) Lung pathology measurements in SARS-CoV-2 MA10 challenged mice from the different groups. Macroscopic lung discoloration score, microscopic acute lung injury (ALI) score, and diffuse alveolar damage (DAD) in day 4 post infection lung tissues are shown. Statistical significance is reported from a one-way ANOVA after Dunnet’s multiple comparison correction. *p < 0.05, **p < 0.01, ***p < 0.001, and ****p < 0.0001.

### Lung pathology and cytokines in mRNA-LNP vaccinated mice challenged with epidemic and pandemic coronaviruses

Lung discoloration is the gross manifestation of various processes of acute lung damage, including congestion, edema, hyperemia, inflammation, and protein exudation. We used this macroscopic scoring scheme to visually score mouse lungs at the time of harvest. To quantify the pathological features of acute lung injury (ALI) in mice, we used a tool from the American Thoracic Society (ATS - Matute-Bello lung pathology score). With a complementary histological quantitation tool, we similarly scored lung tissue sections for diffuse alveolar damage (DAD), the pathological hallmark of ALI (cellular sloughing, necrosis, hyaline membranes, etc.) (*30, 31*) and found these data were consistent with those from the lung discoloration scores. We observed significant lung pathology by both the Matute-Bello and DAD scoring tools in groups 4 and 5 vaccinated animals, consistent with the weight loss and lung titer after heterologous SARS-CoV MA15 challenge. In contrast, multiplexed chimeric spike vaccine formulations in groups 1 and 2 provided complete protection from lung pathology after SARS-CoV MA15 challenge (Fig. 5A and 5B). Mice immunized with the SARS-CoV-2 mRNA vaccine that showed breakthrough infection with SARS-CoV MA15 developed similar lung inflammation as control vaccinated animals, potentially suggesting that future outbreaks of SARS-CoV may cause disease even in individuals vaccinated with SARS-CoV-2. As eosinophilic infiltrates have been observed in vaccinated, 2003 SARS-CoV challenged mice previously (*32*), we analyzed lung tissues in protected vs. infected animals with SARS-CoV MA15 for eosinophilic infiltrates by immunohistochemistry (Fig. S6). Groups 1 and 2 contained rare, scattered eosinophils in the interstitium. Group 3 showed bronchus-associated lymphoid tissue. In contrast, group 4 and group 5 contained frequent perivascular cuffs with prevalent eosinophils. In contrast to the heterologous SARS-CoV MA15 challenge, all groups challenged with SARS-CoV-2 MA10 were protected against lung pathology compared to the norovirus capsid-immunized control group, supporting the hypothesis that the SARS-CoV-2 NTD present in the chimeric spike from group 3 is sufficient for protection (Fig 5C and 5D).

We measured lung proinflammatory cytokines and chemokines in the different vaccination groups. Groups 1 and 2 had baseline levels of macrophage activating cytokines and chemokines including, IL-6, CCL2, IL-1α, G-SCF, and CCL4, compared to group 5 following SARS-CoV MA15 challenge (Fig. S7A). In contrast, group 3 and group 4 showed high and indistinguishable levels of IL-6, CCL2, IL-1α, G-SCF, and CCL4 compared to group 5 mice following SARS-CoV MA15 challenge. Following SARS-CoV-2 MA10 challenge, group 4 and group 1 showed the lowest levels of IL-6, and G-SCF relative to group 5 controls (Fig. S7B), and we only observed significant reductions in CCL2, IL-1α, CCL4 lung levels in groups 3 and 4 compared to the group 5 control despite full protection from both weight loss and lower airway viral replication.

## Discussion

The Moderna and Pfizer/BioNTech SARS-CoV-2 mRNA-LNP vaccines are safe and efficacious against SARS-CoV-2 infections in large Phase 3 efficacy human clinical trials (*33–35*), but there is a growing concern that VOCs like South African B1.351, which is 5-6 fold more resistant to vaccine-elicited polyclonal neutralizing antibodies (*36*). We sought to replicate this platform to establish the breadth of existing SARS-CoV-2 mRNA vaccines, but also to formulate chimeric vaccines that specifically target distant Sarbecovirus strains. A caveat of including multiple chimeric spikes in a single shot is the potential formation of heterotrimers not present in the intended vaccine formulation. While it remains unknown if our chimeric mRNA-LNP vaccines generate heterotrimers *in vivo*, the observation that NTD and RBD chimeric spikes can protect against more than one virus is important. Chimera 4, which contains the RsSHC014 RBD and SARS-CoV-2 NTD and S2, elicited binding and neutralizing antibodies and mice were fully protected from Bt-CoV RsSHC014 and SARS-CoV-2 challenge, whereas SARS-CoV-2 full length did not fully protect against RsSHC014, suggesting that CoV spikes vaccines can be designed to maximize their display of protective epitopes. The lack of protection against WIV-1, SARS-CoV, and only partial protection against RsSHC014 challenge in SARS-CoV-2 immunized mice underlines the need for the development of universal vaccination strategies that can achieve broader coverage against pre-emergent bat SARS-CoV-like and SARS-CoV-2-like viruses. While other strategies exist, including multiplexing mosaic Sarbecovirus RBDs (*37*), RBDs on nanoparticles (*38*), chimeric spike mRNA-LNP vaccination can achieve broad protection using existing manufacturing technologies, and are portable to other high-risk emerging coronaviruses like group 2C MERS-CoV-related strains.

As previously reported with RNA recombinant viruses, our chimeric spike live viruses containing SARS-CoV-2 antigenic domains reaffirm the known interchangeability and functional plasticity of CoV spike glycoprotein structural motifs (*26, 39, 40*). Our demonstration of cross-protection against multiple Sarbecovirus strains in mice, coupled with the absence of immune pathology, lends support to the notion that universal vaccines against group 2B CoVs is likely achievable. Moving forward, it will be important to determine if other combinations of chimeric mRNA-LNP vaccines from other SARS-like viruses are protective, elicit broad T cell responses, prevent the rapid emergence of escape viruses, and elicit protective responses in non-human primate models of Sarbecovirus pathogenesis.

## Author contributions

Conceived the study: D.R.M. and R.S.B. designed experiments: D.R.M, R.S.B., performed laboratory experiments: D.R.M, A.S., S.R.L., A.W.; Provided critical reagents: N.P., K.O.S. Analyzed data and provided critical insight: D.R.M, A.S., S.R.L., G.D.l.C., A.W., E.A.V., L.C.L, N.P., R.P., M.B, D.L., B.Y., K.O.S., D.W., B.F.H., S.M.; Wrote the first draft of the paper: D.R.M; Read and edited the paper: D.R.M, A.S., S.R.L., G.D.l.C., A.W., E.A.V., L.C.L, N.P., N.P., R.P., M.B, D.L., B.Y., K.O.S., D.W., B.F.H., S.M., R.S.B. Funding acquisition: D.R.M., R.S.B. All authors reviewed and approved the manuscript.

## Data and materials availability

The amino acid sequences of the chimeric spike constructs are included in table S1.

## Supplementary materials

### Materials and Methods

#### Chimeric spike vaccine design and formulation

Chimeric spike vaccines were designed with RBD and NTD swaps to increase coverage of epidemic (SARS-CoV), pandemic (SARS-CoV-2), and high-risk pre-emergent bat CoVs (bat SARS-like HKU3-1, and bat SARS-like RsSHC014). Chimeric and monovalent spike mRNA-LNP vaccines were designed based on SARS-CoV-2 spike (S) protein sequence (Wuhan-Hu-1, GenBank: MN908947.3), SARS-CoV (urbani GenBank: AY278741), bat SARS-like CoV HKU3-1 (GenBank: DQ022305), and Bat SARS-like RsSHC014 (GenBank: KC881005). Coding sequences of full-length SARS-CoV-2 furin knockout (RRAR furin cleavage site abolished between amino acids 682-685), the four chimeric spikes, and the norovirus capsid negative control were codon-optimized, synthesized and cloned into the mRNA production plasmid mRNAs were encapsulated with LNP (*41*). Briefly, mRNAs were transcribed to contain 101 nucleotide-long poly(A) tails. mRNAs were modified with m1Ψ-5′-triphosphate (TriLink #N-1081) instead of UTP and the *in vitro* transcribed mRNAs capped using the trinucleotide cap1 analog, CleanCap (TriLink #N-7413). mRNA was purified by cellulose (Sigma-Aldrich # 11363-250G) purification. All mRNAs were analyzed by agarose gel electrophoresis and were stored at −20°C. Cellulose-purified m1Ψ-containing RNAs were encapsulated in proprietary LNPs containing adjuvant (Acuitas) using a self-assembly process as previously described wherein an ethanolic lipid mixture of ionizable cationic lipid, phosphatidylcholine, cholesterol and polyethylene glycol-lipid was rapidly mixed with an aqueous solution containing mRNA at acidic pH. The RNA-loaded particles were characterized and subsequently stored at −80°C at a concentration of 1 mg/ml. The mean hydrodynamic diameter of these mRNA-LNP was ~80 nm with a polydispersity index of 0.02-0.06 and an encapsulation efficiency of ~95%.

#### Animals, immunizations, and challenge viruses

Eleven-month-old female BALB/c mice were purchased from Envigo (#047) and were used for all experiments. The study was carried out in accordance with the recommendations for care and use of animals by the Office of Laboratory Animal Welfare (OLAW), National Institutes of Health and the Institutional Animal Care and Use Committee (IACUC) of University of North Carolina (UNC permit no. A-3410-01). mRNA-LNP vaccines were kept frozen until right before the vaccination. Mice were immunized with a total 1μg in the prime and boost. Briefly, chimeric vaccines were mixed at 1:1 ratio for a total of 1μg when more than one chimeric spike was used or 1μg of a single spike diluted in sterile 1XPBS in a 50μl volume and were given 25μl intramuscularly in each hind leg. Prime and boost immunizations were given three weeks apart. Three weeks post boost, mice were bled, sera was collected for analysis, and mice were moved into the BSL3 facility for challenge experiments. Animals were housed in groups of five and fed standard chow diets. Virus inoculations were performed under anesthesia and all efforts were made to minimize animal suffering. All mice were anesthetized and infected intranasally with 1 × 10^4^ PFU/ml of SARS-CoV MA15, 1 × 10^4^ PFU/ml of SARS-CoV-2 MA10, 1 × 10^4^ PFU/ml RsSHC014, 1 × 10^4^ PFU/ml RsSHC014-MA15, 1 × 10^5^ PFU/ml WIV-1, and 1 × 10^4^ PFU/ml SARS-CoV-2 B.1.351-MA10 which have been described previously (*8, 42, 43*). Mice were weighted daily and monitored for signs of clinical disease. Each challenge virus challenge experiment encompassed 50 mice with 10 mice per vaccine group to obtain statistical power. Mouse vaccinations and challenge experiments were independently repeated twice to ensure reproducibility.

#### Measurement of mouse CoV spike binding antibodies by ELISA

Mouse serum samples from pre-immunization (pre-prime), 2 weeks post prime (pre-boost), and 3 weeks post boost were tested. A binding ELISA panel that included SARS-CoV spike Protein DeltaTM, SARS-CoV-2 (2019-nCoV) spike Protein (S1+S2 ECD, His tag), MERS-CoV, Coronavirus spike S1+S2 (Baculovirus-Insect Cells, His), HKU1 (isolate N5) spike Protein (S1+S2 ECD, His Tag), OC43 spike Protein (S1+S2 ECD, His Tag), 229E spike Protein (S1+S2 ECD, His tag) Human coronavirus (HCoV-NL63) spike Protein (S1+S2 ECD, His Tag), Pangolin CoV_GXP4L_spikeEcto2P_3C8HtS2/293F, bat CoV RsSHC014_spikeEcto2P_3C8HtS2/293F, RaTG13_spikeEcto2P_3C8HtS2/293F, and bat CoV HKU3-1 spike were tested. Indirect binding ELISAs were conducted in 384 well ELISA plates (Costar #3700) coated with 2μg/ml antigen in 0.1M sodium bicarbonate overnight at 4°C, washed and blocked with assay diluent (1XPBS containing 4% (w/v) whey protein/ 15% Normal Goat Serum/ 0.5% Tween-20/ 0.05% Sodium Azide). Serum samples were incubated for 60 minutes in three-fold serial dilutions beginning at 1:30 followed by washing with PBS/0.1% Tween-20. HRP conjugated goat anti-mouse IgG secondary antibody (SouthernBiotech 1030-05) was diluted to 1:10,000 in assay diluent without azide, incubated at for 1 hour at room temperature, washed and detected with 20µl SureBlue Reserve (KPL 53-00-03) for 15 minutes. Reactions were stopped via the addition of 20µl HCL stop solution. Plates were read at 450nm. Area under the curve (AUC) measurements were determined from binding of serial dilutions.

#### ACE2 blocking ELISAs

Plates were coated with 2μg/ml recombinant ACE2 protein, then washed and blocked with 3% BSA in PBS. While assay plates blocked, and sera was diluted 1:25 in 1%BSA/0.05% Tween-20. Then SARS-CoV-2 spike protein was mixed with equal volumes of each sample at a final spike concentration equal to the EC_50_ at which it binds to ACE2. The mixture was allowed to incubate at room temperature for 1 hour. Blocked assay plates were washed, and the serum-spike mixture was added to the assay plates for a period of 1 hour at room temperature. Plates were washed and Strep-Tactin HRP, (IBA GmbH, Cat# 2-1502-001) was added at a dilution of 1:5000 followed by TMB substrate. The extent to which antibodies were able to block the binding of spike protein to ACE2 was determined by comparing the OD of antibody samples at 450nm to the OD of samples containing spike protein only with no antibody. The following formula was used to calculate percent blocking (100-(OD sample/OD of spike only) *100).

#### Measurement of neutralizing antibodies against live viruses

Full-length SARS-CoV-2 Seattle, SARS-CoV-2 D614G, SARS-CoV-2 B.1.351, SARS-CoV-2 B.1.1.7, SARS-CoV-2 mink, SARS-CoV, WIV-1, and RsSHC014 viruses were designed to express nanoluciferase (nLuc) and were recovered via reverse genetics as described previously (*29*). Virus titers were measured in Vero E6 USAMRIID cells, as defined by plaque forming units (PFU) per ml, in a 6-well plate format in quadruplicate biological replicates for accuracy.

For the 96-well neutralization assay, Vero E6 USAMRID cells were plated at 20,000 cells per well the day prior in clear bottom black walled plates. Cells were inspected to ensure confluency on the day of assay. Serum samples were tested at a starting dilution of 1:20 and were serially diluted 3-fold up to nine dilution spots. Serially diluted serum samples were mixed in equal volume with diluted virus. Antibody-virus and virus only mixtures were then incubated at 37°C with 5% CO_2_ for one hour. Following incubation, serially diluted sera and virus only controls were added in duplicate to the cells at 75 PFU at 37°C with 5% CO_2_. After 24 hours, cells were lysed, and luciferase activity was measured via Nano-Glo Luciferase Assay System (Promega) according to the manufacturer specifications. Luminescence was measured by a Spectramax M3 plate reader (Molecular Devices, San Jose, CA). Virus neutralization titers were defined as the sample dilution at which a 50% reduction in RLU was observed relative to the average of the virus control wells.

#### Eosinophilic lung infiltrates staining

To detect eosinophils, chromogenic immunohistochemistry (IHC) was performed on paraffin-embedded lung tissues that were sectioned at 4 microns. Lung tissues from vaccine groups 1-5 were analyzed for lung eosinophilic infiltration. N=8-10 lung tissues per group were analyzed. This IHC was carried out using the Leica Bond III Autostainer system. Slides were dewaxed in Bond Dewax solution (AR9222) and hydrated in Bond Wash solution (AR9590). Heat induced antigen retrieval was performed for 20 min at 100°C in Bond-Epitope Retrieval solution 2, pH-9.0 (AR9640). After pretreatment, slides were incubated with an Eosinophil Peroxidase antibody (PA5-62200, Invitrogen) at 1:1,000 for 1h followed with Novolink Polymer (RE7260-K) secondary. Antibody detection with 3,3’-diaminobenzidine (DAB) was performed using the Bond Intense R detection system (DS9263). Stained slides were dehydrated and coverslipped with Cytoseal 60 (8310-4, Thermo Fisher Scientific). Two positive controls (one with high and another with low eosinophil reactivity) and a negative control (no primary antibody) were included in all staining runs.

#### Lung pathology scoring

Acute lung injury was quantified via two separate lung pathology scoring scales: Matute-Bello and Diffuse Alveolar Damage (DAD) scoring systems. Analyses and scoring were performed by a board vertified veterinary pathologist who was blinded to the treatment groups as described previously (*30*). Lung pathology slides were read and scored at 600X total magnification.

The lung injury scoring system used is from the American Thoracic Society (Matute-Bello) in order to help quantitate histological features of ALI observed in mouse models to relate this injury to human settings. In a blinded manner, three random fields of lung tissue were chosen and scored for the following: (A) neutrophils in the alveolar space (none = 0, 1–5 cells = 1, > 5 cells = 2), (B) neutrophils in the interstitial septa (none = 0, 1–5 cells = 1, > 5 cells = 2), (C) hyaline membranes (none = 0, one membrane = 1, > 1 membrane = 2), (D) Proteinaceous debris in air spaces (none = 0, one instance = 1, > 1 instance = 2), (E) alveolar septal thickening (< 2x mock thickness = 0, 2–4x mock thickness = 1, > 4x mock thickness = 2). To obtain a lung injury score per field, A–E scores were put into the following formula score = [(20x A) + (14 x B) + (7 x C) + (7 x D) + (2 x E)]/100. This formula contains multipliers that assign varying levels of importance for each phenotype of the disease state. The scores for the three fields per mouse were averaged to obtain a final score ranging from 0 to and including 1.

The second histology scoring scale to quantify acute lung injury was adopted from a lung pathology scoring system from lung RSV infection in mice (*31*). This lung histology scoring scale measures diffuse alveolar damage (DAD). Similar to the implementation of the ATS histology scoring scale, three random fields of lung tissue were scored for the following in a blinded manner: 1= absence of cellular sloughing and necrosis, 2=Uncommon solitary cell sloughing and necrosis (1–2 foci/field), 3=multifocal (3+foci) cellular sloughing and necrosis with uncommon septal wall hyalinization, or 4=multifocal (>75% of field) cellular sloughing and necrosis with common and/or prominent hyaline membranes. The scores for the three fields per mouse were averaged to get a final DAD score per mouse. The microscope images were generated using an Olympus Bx43 light microscope and CellSense Entry v3.1 software.

#### Measurement of lung cytokines

Lung tissue was homogenized, spun down at 13,000g, and supernantant was used to measure lung cytokines using Mouse Cytokine 23-plex Assay (BioRad). Briefly, 50μl of lung homogenate supernatant was added to each well and the protocol was followed according to the manufacturer specifications. Plates were read using a MAGPIX multiplex reader (Luminex Corporation).

#### Biocontainment and biosafety

Studies were approved by the UNC Institutional Biosafety Committee approved by animal and experimental protocols in the Baric laboratory. All work described here was performed with approved standard operating procedures for SARS-CoV-2 in a biosafety level 3 (BSL-3) facility conforming to requirements recommended in the Microbiological and Biomedical Laboratories, by the U.S. Department of Health and Human Service, the U.S. Public Health Service, and the U.S. Center for Disease Control and Prevention (CDC), and the National Institutes of Health (NIH).

#### Statistics

All statistical analyses were performed using GraphPad Prism 9. Statistical tests used in each figure are denoted in the corresponding figure legend.

---

**Figure S1.**
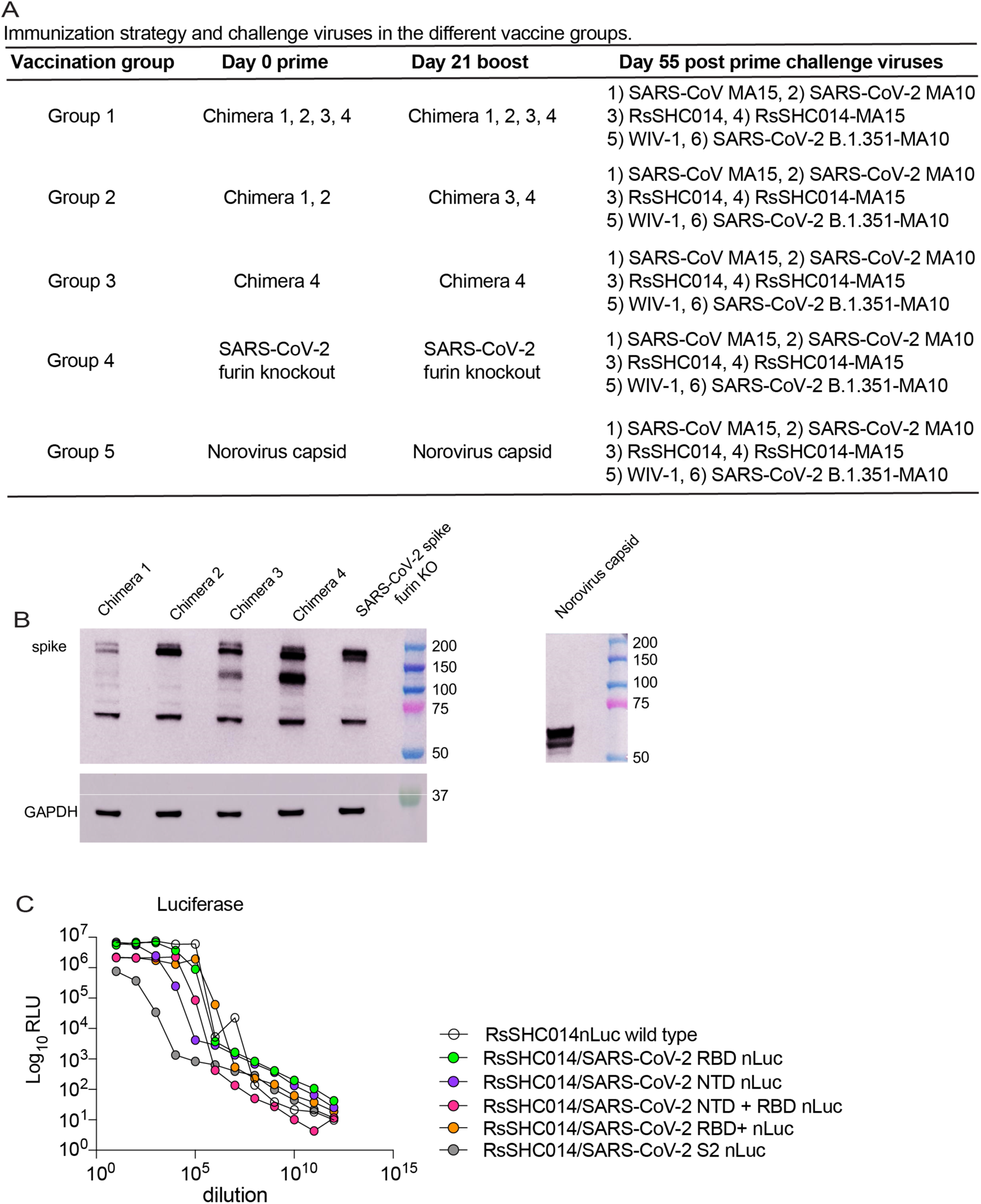
Chimeric and wild type spike Sarbecovirus constructs. (**A**) Mouse vaccination strategy using mRNA-LNPs: group 1 received chimeric spike 1, 2, 3, and 4 as the prime and boost, group 2 received chimeric spike 1, 2 as the prime and chimeric spikes 3 and 4 as the boost, group 3 received chimeric spike 4 as the prime and boost, group 4 received SARS-CoV-2 furin KO prime and boost, and group 5 received a norovirus capsid prime and boost. Different vaccine groups were separately challenged with 1) SARS-CoV MA15, 2) SARS-CoV-2 MA10, 3) RsSHC014 full-length virus, 4) RsSHC014-MA15, 5) WIV-1, and 6) SARS-CoV-2 B.1.351 MA10. (**B**) Protein expression of chimeric spikes, SARS-CoV-2 furin KO, and norovirus mRNA vaccines. The extra band between 100-150 kDa corresponds to S1. GAPDH was used as the loading control. (**C**) Nanoluciferase expression of RsSHC014/SARS-CoV-2 chimeric spike live viruses.

**Figure S2.**
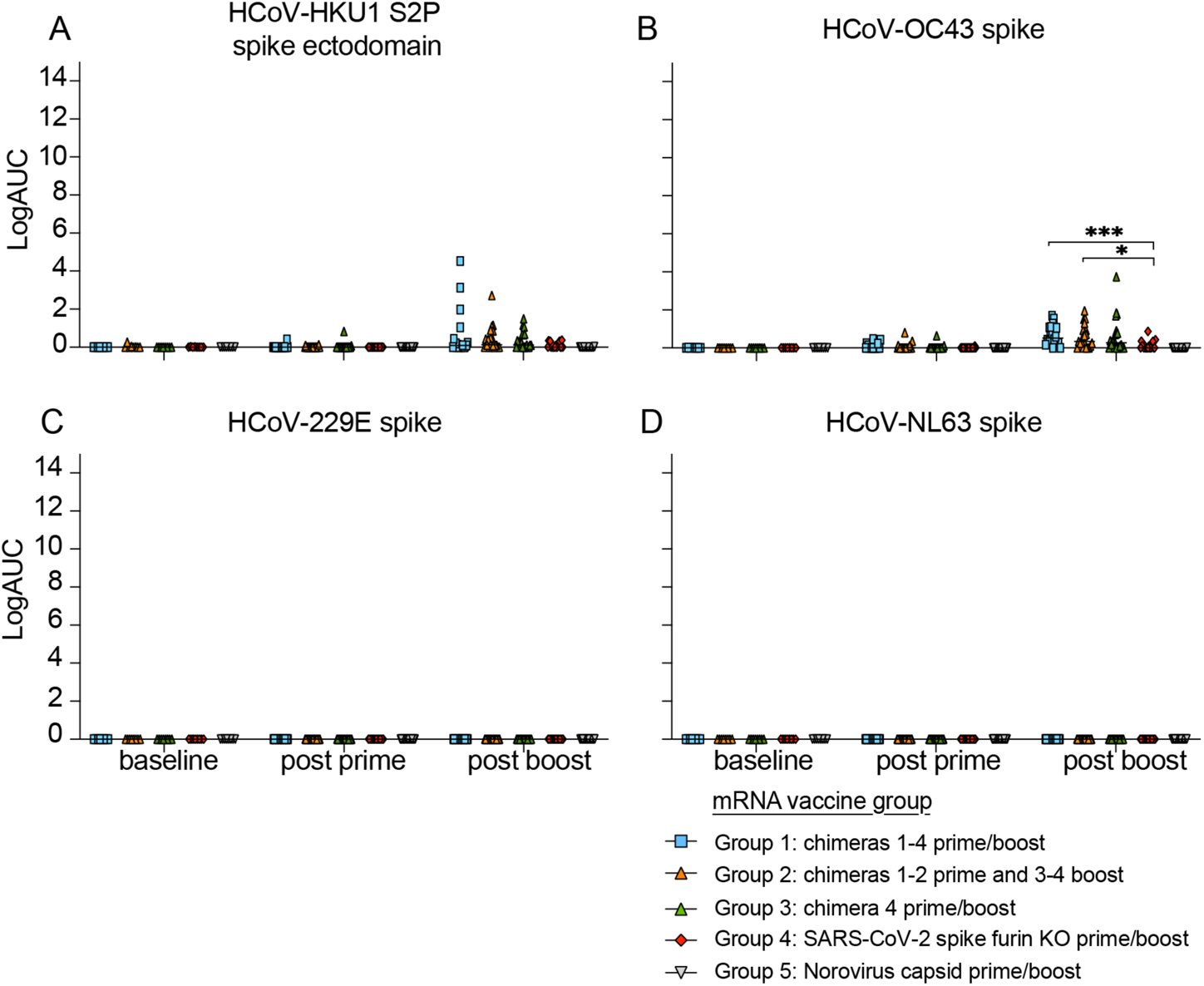
Human common-cold CoV ELISA binding responses in chimeric and monovalent SARS-CoV-2 spike mRNA-LNP-vaccinated mice. Pre-immunization, post prime, and post boost binding to (**A**) HCoV-HKU1 spike, (**B**) HCoV-OC43 spike, (**C**) HCoV-229E spike, and (**D**) HCoV-NL63 spike. Statistical significance for the binding and blocking responses is reported from a Kruskal-Wallis test after Dunnett’s multiple comparison correction. *p < 0.05, **p < 0.01, ***p < 0.001, and ****p < 0.0001.

**Figure S3.**
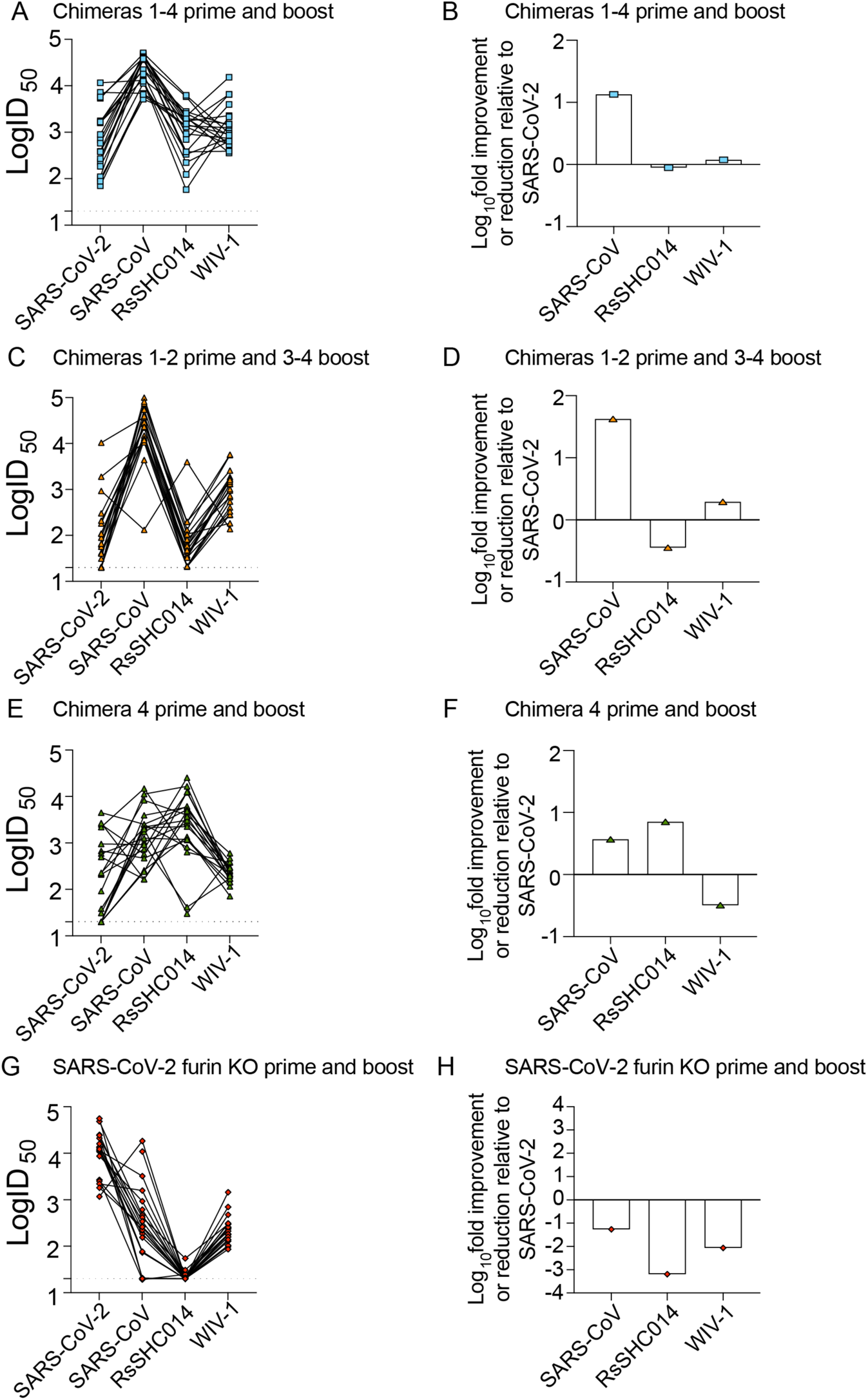
Comparison of neutralizing antibody activity of CoV mRNA-LNP vaccines against Sarbecoviruses. (**A**) Group 1 neutralizing antibody responses against SARS-CoV-2, SARS-CoV, RsSHC014, and WIV-1 and (**B**) fold-change of SARS-CoV, RsSHC014, and WIV-1 neutralizing antibodies relative to SARS-CoV-2. (**C**) Group 2 neutralizing antibody responses against SARS-CoV-2, SARS-CoV, RsSHC014, and WIV-1 and (**D**) fold-change of SARS-CoV, RsSHC014, and WIV-1 neutralizing antibodies relative to SARS-CoV-2. (**E**) Group 3 neutralizing antibody responses against SARS-CoV-2, SARS-CoV, RsSHC014, and WIV-1 and (**F**) fold-change of SARS-CoV, RsSHC014, and WIV-1 neutralizing antibodies relative to SARS-CoV-2. (**G**) Group 4 neutralizing antibody responses against SARS-CoV-2, SARS-CoV, RsSHC014, and WIV-1 and (**H**) fold-change of SARS-CoV, RsSHC014, and WIV-1 neutralizing antibodies relative to SARS-CoV-2.

**Figure S4.**
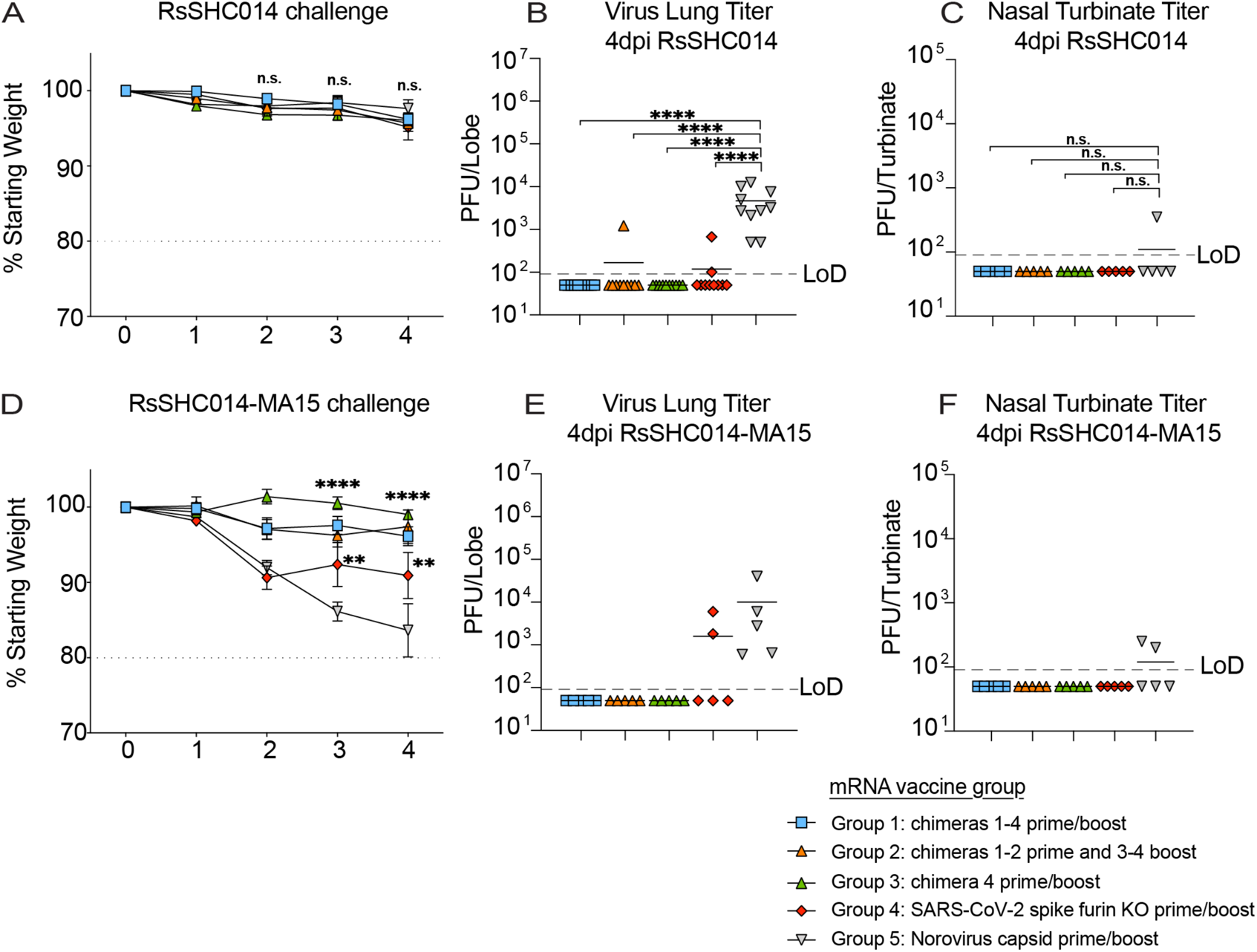
*In vivo* protection against Bt-CoV challenge by chimeric spikes mRNA-vaccines. (**A**) Percent starting weight from the different vaccine groups of mice challenged with full-length RsSHC014. (**B**) RsSHC014 lung viral titers in mice from the distinct vaccine groups. (**C**) RsSHC014 nasal turbinate titers in mice from the different immunization groups. (**D**) Percent starting weight from the different vaccine groups of mice challenged with RsSHC014-MA15. (**E**) RsSHC014-MA15 lung viral titers in mice from the distinct vaccine groups. (**F**) RsSHC014-MA15 nasal turbinate titers in mice from the different immunization groups. Statistical significance is reported from a one-way ANOVA after Tukey’s multiple comparison correction. *p < 0.05, **p < 0.01, ***p < 0.001, and ****p < 0.0001.

**Figure S5.**
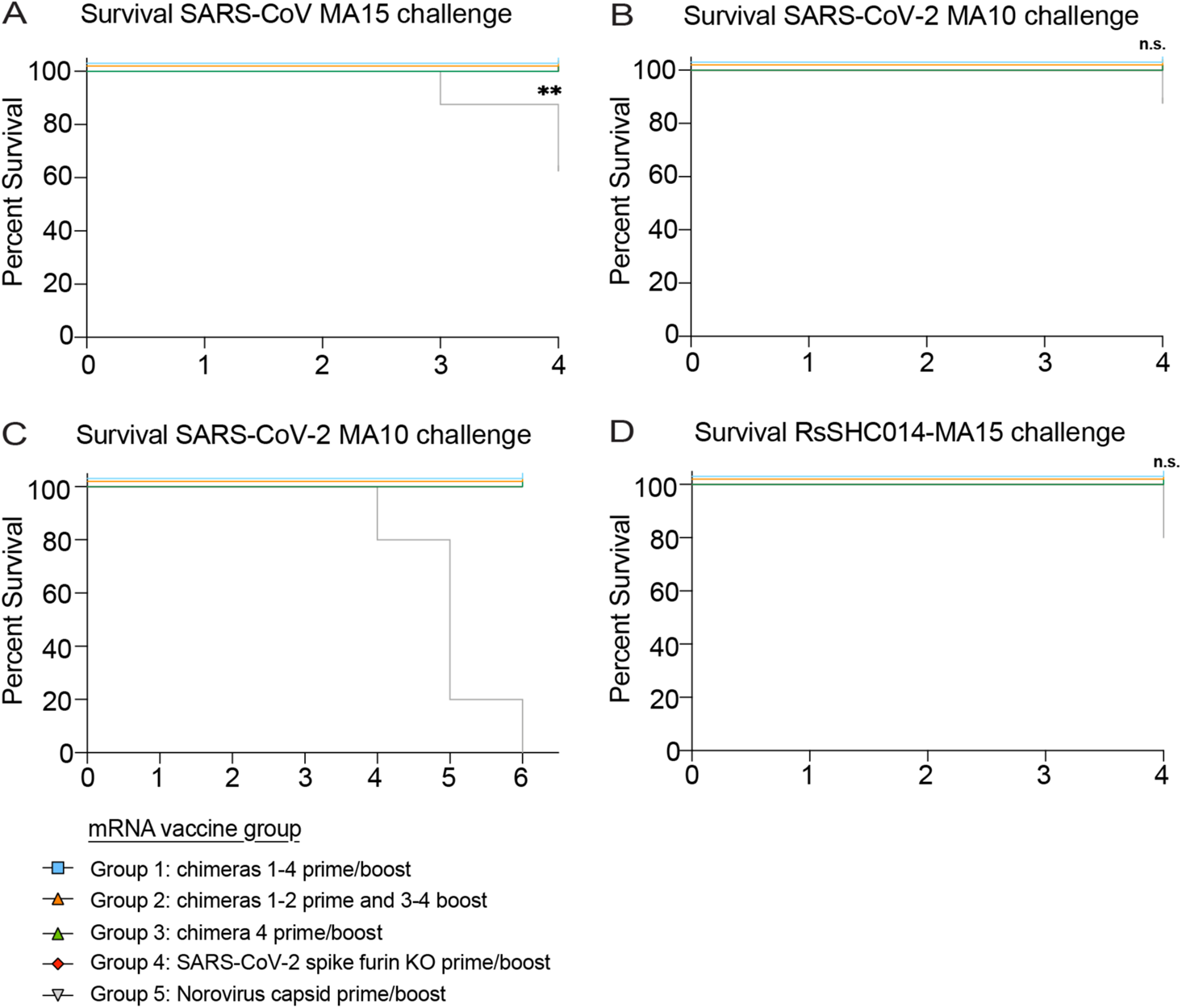
Survival analysis of immunized mice challenged with Sarbecoviruses. (**A**) Survival analysis at day 4 post infection from immunized mice infected with SARS-CoV MA15, (**B**) SARS-CoV-2 MA10, (**C**) Survival analysis at day 7 post infection from immunized mice infected with SARS-CoV-2 MA10, and (**D**) RsSHC014-MA15. Statistical significance is reported from a Mantel-Cox test.

**Figure S6.**
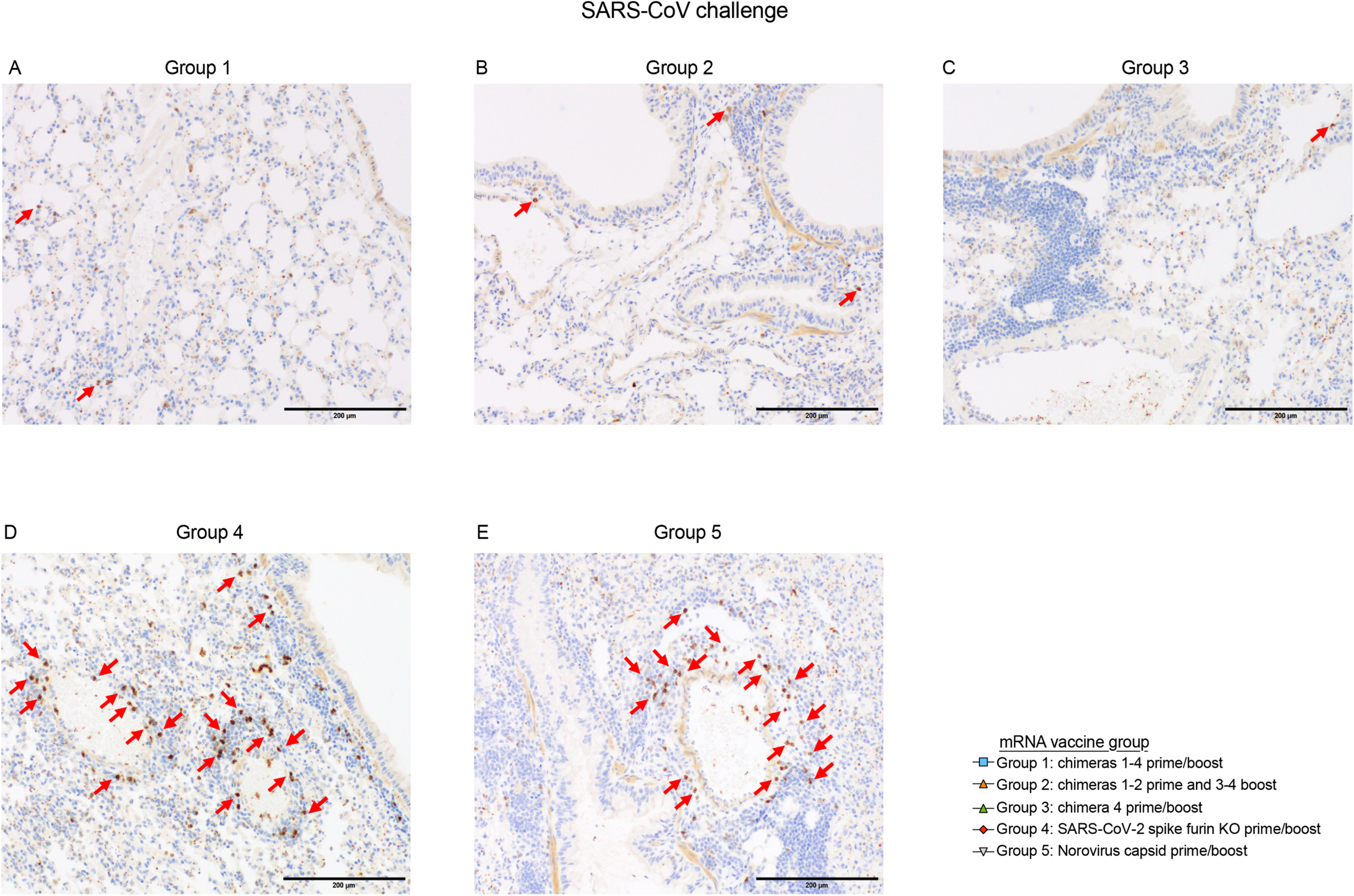
Detection of eosinophilic infiltrates in SARS-CoV MA15 challenged mice. (**A**) Group 1: rare scattered individual eosinophils in the interstitium with some small perivascular cuffs that lack eosinophils. (**B**) Group 2: Bronchiolar cuffs of leukocytes with rare eosinophils. (**C**) Group 3: Hyperplastic bronchus-associated lymphoid tissue (BALT) with rare eosinophils. (**D**) Group 4: frequent perivascular cuffs that contain eosinophils. (**E**) Group 5: frequent eosinophils in perivascular cuffs.

**Figure S7.**
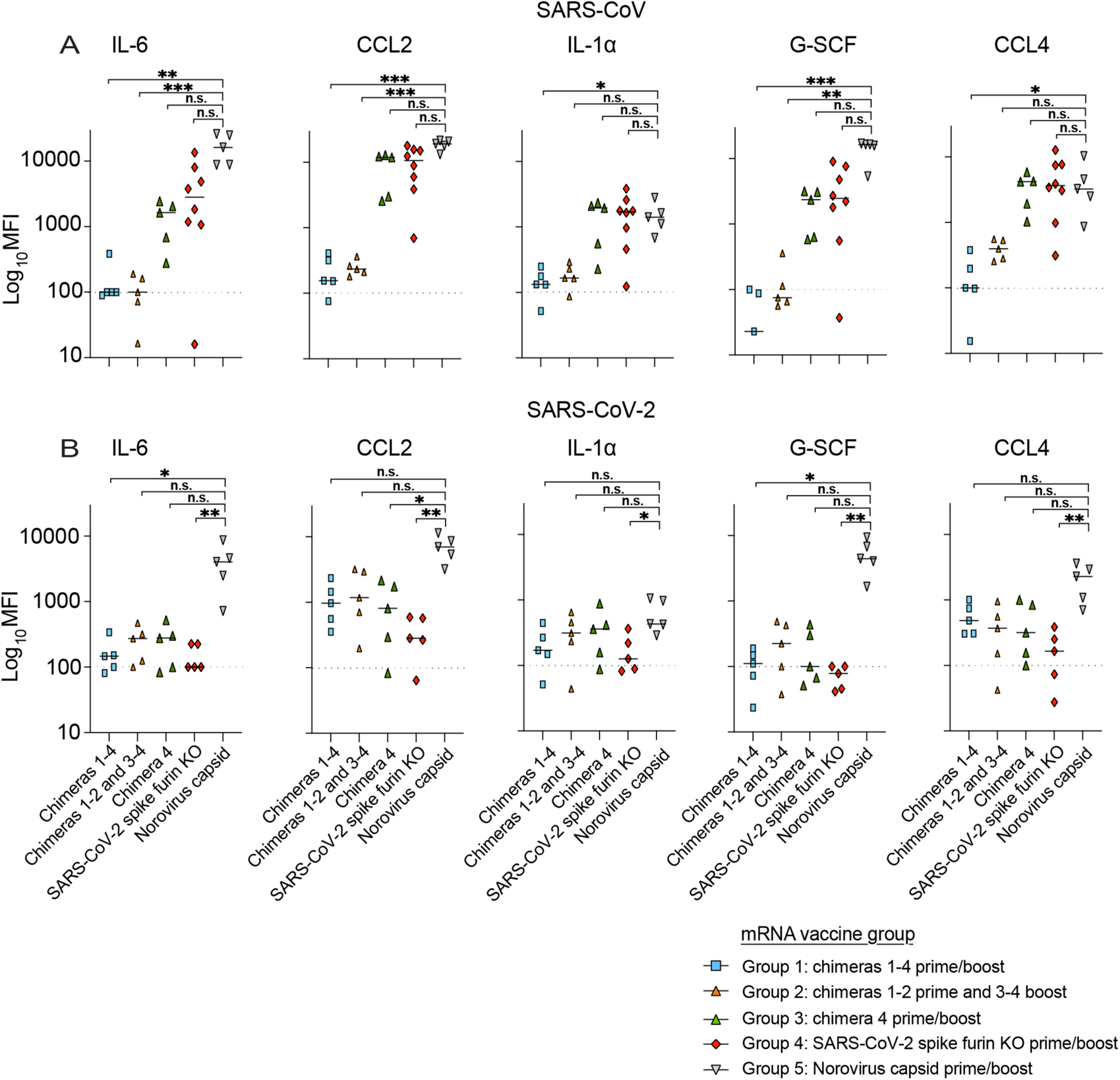
Lung cytokine analysis in Sarbecovirus-challenged mice. CCL2, IL-1α, G-SCF, and CCL4 in (**A**) SARS-CoV-infected mice and in (**B**) SARS-CoV-2-infected mice. Statistical significance for the binding and blocking responses is reported from a Kruskal-Wallis test after Dunnett’s multiple comparison correction. *p < 0.05, **p < 0.01, ***p < 0.001, and ****p < 0.0001.

**Table S1:**
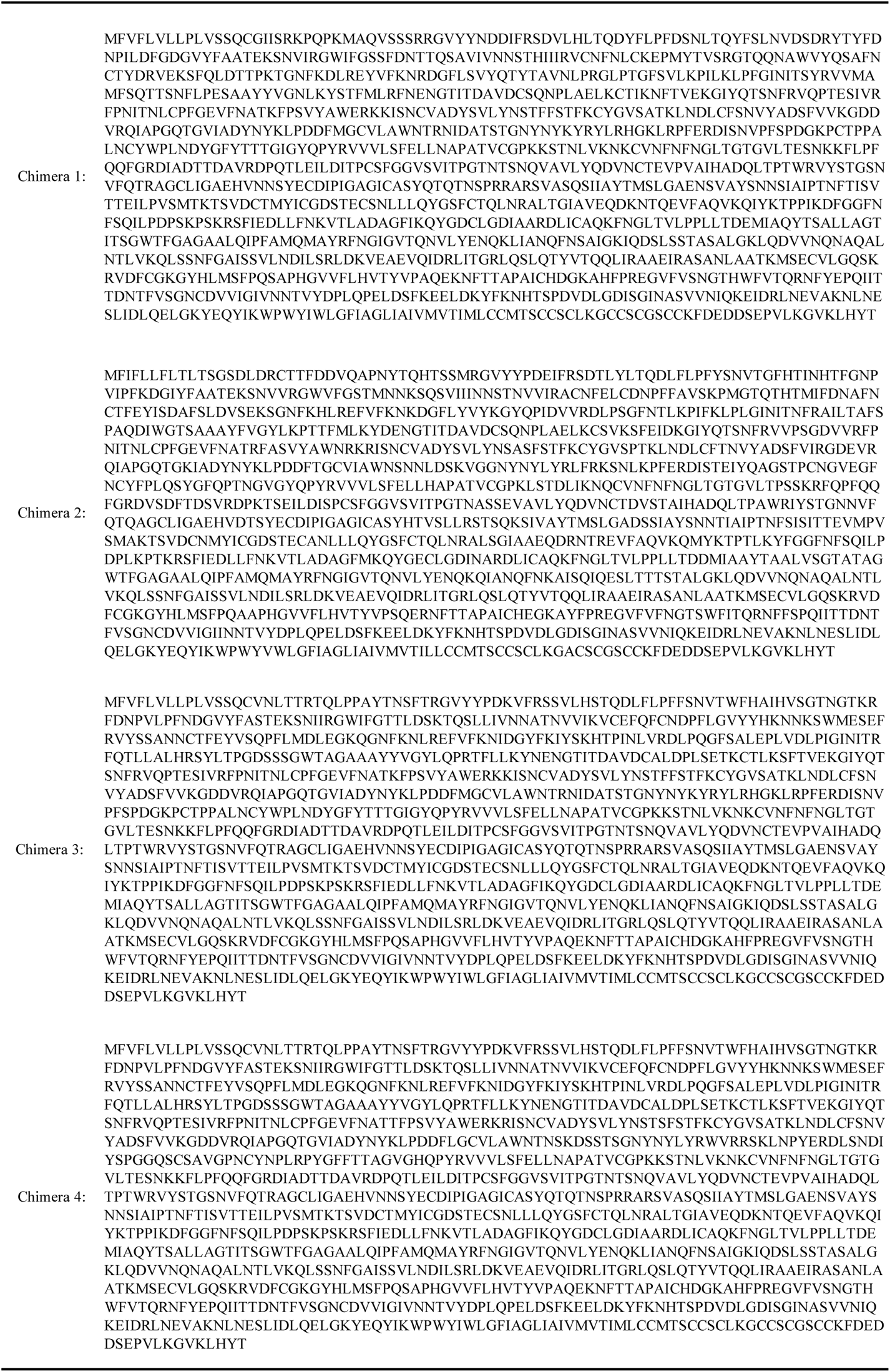
Amino acid sequences of chimeric spikes.

